# Neural representations of concrete concepts enable identification of individuals during naturalistic story listening

**DOI:** 10.1101/2023.09.07.556725

**Authors:** Thomas L. Botch, Emily S. Finn

**Affiliations:** Department of Psychological & Brain Sciences, Dartmouth College, Hanover, NH, 03755

**Author notes:** Corresponding author: Thomas L. Botch.

## Abstract

Different people listening to the same story may converge upon a largely shared interpretation while still developing idiosyncratic experiences atop that shared foundation. What semantic properties support this individualized experience of natural language? Here, we investigate how the “concreteness” of word meanings — i.e., the extent to which a concept is derived from sensory experience — relates to variability in the neural representations of language. Leveraging a large dataset of participants who each listened to four auditory stories while undergoing functional MRI, we demonstrate that an individual’s neural representations of concrete concepts are reliable across stories and unique to the individual. In contrast, we find that neural representations of abstract concepts are variable both within individuals and across the population. Using natural language processing tools, we show that concrete words exhibit similar neural signatures despite spanning larger distances within a high-dimensional semantic space, which potentially reflects an underlying signature of sensory experience — namely, imageability — shared by concrete words but absent from abstract words. Our findings situate the concrete-abstract semantic axis as a core dimension that supports reliable yet individualized representations of natural language.

## Introduction

The success of language as a means of communication relies on a shared understanding of the meanings of words as links to mental concepts^1–3^. While there is generally strong convergence in how people understand and represent language^4,5^, the conceptual associations evoked by a given word can also be highly individualized and informed by experience^6,7^. What semantic properties scaffold common conceptual knowledge while also providing the foundation for idiosyncratic representations?

A large body of empirical and theoretical work has suggested that human knowledge is organized along an axis that moves from concrete, sensory-based representations to abstract, language-derived representations^8–11^. These theories propose that concrete concepts benefit from being represented across both sensory and linguistic domains and, as a result, exhibit more stable representations than abstract concepts. Recent findings from human neuroimaging support these theories, demonstrating close topographical and functional correspondence between representations of sensory and linguistic information^12–14^. Furthermore, neural representations of concrete concepts are less variable across subjects than representations of abstract concepts^15–17^. In turn, the stability of concrete concept representations is suggested to benefit behavior: concrete words are processed faster^16,18–20^, are more imageable^21,22^, and are more easily recalled than abstract words^23–27^. While these studies suggest population-level commonalities in how people process and represent the concrete-abstract axis, the extent to which these representations are colored by individual experience remains unclear.

More recently, language researchers have demonstrated differences in how individuals organize and represent word meanings^17^, finding that concrete words demonstrate greater similarity across subjects than abstract words, in both conceptual organization (as measured behaviorally with a semantic distance task) and neural representation. This result further suggests that representations become more consistent across subjects as words become more concrete and more variable as words become more abstract. However, the low similarity of abstract word representations across subjects could stem from multiple causes (Figure 1C). On one hand, representations of abstract words might be highly individualized—in other words, unique and colored by an individual’s experiences. Such individual-specific representations would be evidenced by high within-subject similarity across repeated exposures to the same word or concept, despite low across-subject similarity. Another possibility is that low similarity results from unstable representations of abstract words. In this case, representations would show low similarity both within and across subjects that could result from high variability in the meaning of abstract words across contexts. Yet, without evaluating the reliability of representations *within* subjects, the low similarity of abstract word representations *across* subjects is difficult to interpret.

**Figure 1.**
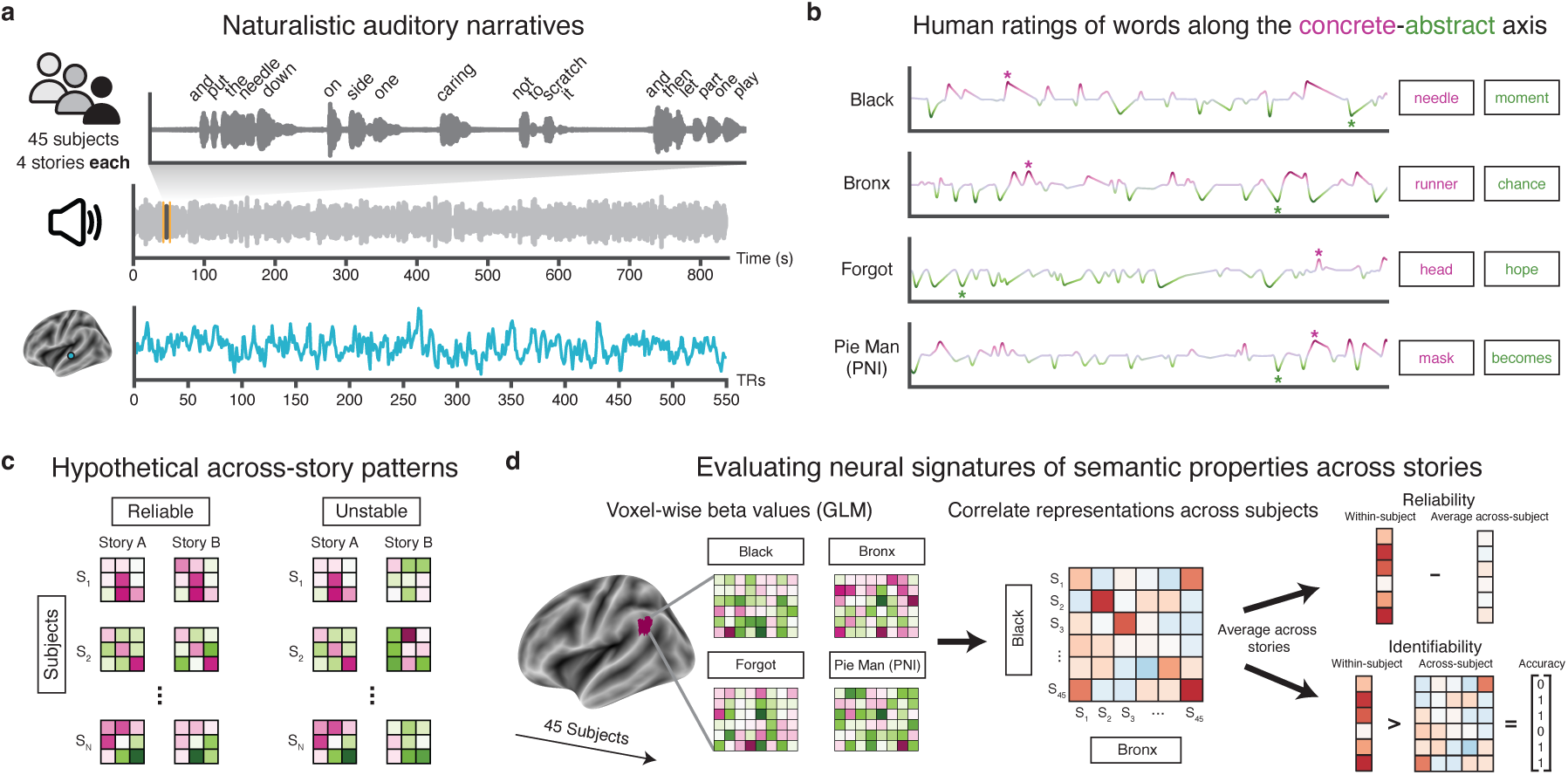
Experimental methods. **(a)** 45 subjects listened to four auditory stories during fMRI scanning (Nastase et al., 2021). **(b)** Human ratings were used to assign a value of concreteness (i.e., position along the concrete-abstract axis) for as many words as possible within each story. This process was repeated with other semantic properties including frequency, valence, and arousal. **(c)** Any apparent variation across subjects in neural representations of linguistic properties could stem from two possible underlying patterns: neural representations could be reliably idiosyncratic within subjects, evidenced by high similarity of representations within the same subject across distinct experiences (here, stories), or these representations could be unstable both within and across subjects, evidenced by variability within the same subject across stories. **(d)** For each story, voxel-wise beta values were estimated for each linguistic property within a generalized linear model. Then, within each of 200 parcels (Schaefer parcellation), beta values were correlated between all subjects for each pair of stories, and story similarity matrices were averaged across all pairs of stories. From these average similarity matrices, we estimate two indices of within-subject stability of neural representations: 1) reliability, defined as the difference between within-subject and average across-subject similarity, and 2) identifiability, defined as the fingerprinting accuracy of discriminating one subject from all other subjects based on their neural representations.

Here, we aimed to understand how the concrete-abstract axis provides a foundation for individual differences in the neural representation of language. We investigated this question within a large dataset of subjects who listened to four naturalistic auditory stories during functional magnetic resonance imaging (fMRI) scanning. Unlike many previous investigations that used isolated single-word or otherwise simplified paradigms^15–17,28–32^, these data allowed us to characterize neural representations of concrete and abstract words as they occur in contextualized speech, as language is used in everyday life^33^. We tested not only the extent to which neural representations of concrete and abstract words are consistent across a group of subjects, but also the degree to which these representations are reliable within and unique to a given subject across stories. Then, by leveraging tools from natural language processing, we examined how the organization of words within a high-dimensional semantic space promotes differential reliability of neural representations of concrete and abstract words.

## Methods

### Participants

We used data from 45 subjects (N=33 female; mean age 23.3 +/- 7.4 years) from the publicly available *Narratives* dataset^34^ who listened to four auditory stories (“Running from the Bronx”, 8:56 min; “Pie Man (PNI)”, 6:40 min; “I Knew You Were Black”, 13:20 min; “The Man Who Forgot Ray Bradbury”, 13:57 min) during fMRI scans at the Princeton Neuroscience Institute (Figure 1A). All stories were collected within the same testing session and each story was collected within a separate run. Across participants, the order of stories was pseudo-randomized such that “Bronx” and “Pie Man (PNI)” were always presented in the first half of the session while “Black” and “Forgot” were presented in the second half of the session. The order of the stories presented within each half of the session was then randomized, resulting in four possible presentation orders across participants. All participants completed written informed consent, were screened for MRI safety and reported fluency in English, having normal hearing, and no history of neurological disorders. The study was approved by the Princeton University Institutional Review Board.

### MRI data acquisition and preprocessing

Functional and anatomical images were collected on a 3T Siemens Magnetom Prisma with a 64-channel head coil. Whole-brain images were acquired (48 slices per volume, 2.5mm isotropic resolution) in an interleaved fashion using a gradient-echo EPI (repetition time (TR) = 1.5s, echo time (TE) = 31ms, flip angle (FA) = 67°) with a multiband acceleration factor of 3 and no in-plane acceleration. A total of 1717 volumes were collected for each participant across four separate scan runs, where a single story was presented within each run.

We used preprocessed data provided by Nastase et al., 2021. In brief, data were preprocessed using fMRIPrep^35^ including co-registration, slice-time correction, and non-linear alignment to the MNI152 template brain. Time-series were detrended with regressors for motion, white matter, cerebrospinal fluid and smoothed with a 6mm FWHM gaussian kernel. For more information about data acquisition and preprocessing, please refer to Nastase et al., 2021.

As an additional preprocessing step, we performed functional alignment on these data using a shared response model^36^ as implemented in *BrainIAK*^37^. Previous work has demonstrated better functional alignment by fitting a SRM within each parcel^38^. Accordingly, we restricted our analyses to the neocortex and used the 200-parcel, 17-network Schaefer parcellation^39^ and removed any parcel without at least 75% coverage across all participants and stories (total parcels removed: 9/200, or 4.5%). Within each remaining parcel, we then fit a model to capture reliable responses to all stories across participants in a lower dimensional feature space (number of features = 50). We then inverted the parcel-wise models to reconstruct the individual voxel-wise time courses for each participant and each story^40^. This procedure served as an additional denoising step to improve the consistency of stimulus-driven spatiotemporal patterns across participants. All analyses were conducted in volume space and projected to surface space for visualization purposes only.

### Stimulus preprocessing

Each story was originally transcribed and aligned to the audio file using the Gentle forced-alignment algorithm by the authors of Nastase et al., 2021. We applied additional preprocessing to the transcripts using the Natural Language Toolkit^41^. First, we obtained parts-of-speech and word lemmas — the base form of a word (e.g., “go” is the lemma for “going”, “gone” and “went”) — for each word, and excluded stop-words (uninformative, common words) such as “the”, “a”, and “is”.

To address our hypotheses, we leveraged an existing corpus of human ratings of word concreteness^42^. In this study, online participants rated a total of 40,000 English word lemmas on a 5-point Likert scale from abstract (lower) to concrete (higher). Each word was rated by at least 25 participants. Participants were instructed to consider a word as more concrete if it refers to something that exists in reality and can be experienced directly through senses or actions, and, in contrast, to consider a word as more abstract if its meaning is dependent on language and cannot be experienced directly through senses or actions. Henceforth, we use “concrete-abstract axis” to refer to this general semantic dimension, and “concreteness” as a word’s specific position on this axis.

For each word in each story, we assigned a value of concreteness using the average human rating for that word’s lemma if it was present in the concreteness corpus (Figure 1B). In addition to our critical predictor (concreteness), we included three other semantic properties as controls: frequency^43,44^, a measure of how often a word occurs in language, and two affective properties, valence and arousal^45^. Word frequency was derived objectively by calculating the number of occurrences of a word per million words (51 million total words), while valence and arousal were derived from human ratings analogous to the concreteness ratings described above. Previous research investigating word frequency effects have demonstrated that less frequent words drive stronger neural responses within the language network^46,47^. A separate set of studies investigating affect have demonstrated that valence and arousal contribute to representations of language within areas related to emotion processing and memory^48,49^. While the selected control semantic properties are not a definitive list, including them as “competition” allows us to make inferences that are more specific to the concrete-abstract axis. Our analysis was then constrained to the set of words with a value for any of the four properties (i.e., the union), resulting in 93% of content words sampled on average across stories.

### fMRI Analysis

#### Modeling representations of semantic properties

For each story and participant, we used a general linear model (GLM) to estimate BOLD responses for each semantic property (concreteness, frequency, valence, arousal), plus a low-level auditory feature regressor (loudness: the root mean square of the auditory waveform). We collectively refer to these semantic and auditory properties as “linguistic properties”. Specifically, to construct a continuous, amplitude-modulated regressor, each linguistic property was assigned a value at each timepoint of the story timeseries based on the word(s) spoken at that timepoint. We then modeled these linguistic properties using AFNI^50^. The model yields a map of beta values that correspond to responses to each property, where higher and lower values indicate higher and lower values of a given linguistic property (e.g., higher = more concrete, lower = more abstract). As all linguistic properties were included in the same model, the resulting beta values represent the BOLD response to a given property while controlling for all other properties.

Using the outputs from these models, we first examined group-level univariate responses to each linguistic property using a linear-mixed effects model. At each voxel, the model predicts BOLD activity from the fixed effects of each linguistic property plus the random effects of subject and story. The model therefore yields a map of beta values that describes consistent neural responses to each linguistic property across stories and subjects. All voxel-wise results are shown following correction for multiple comparisons (FDR *q* < 0.05; Figure 2).

**Figure 2.**
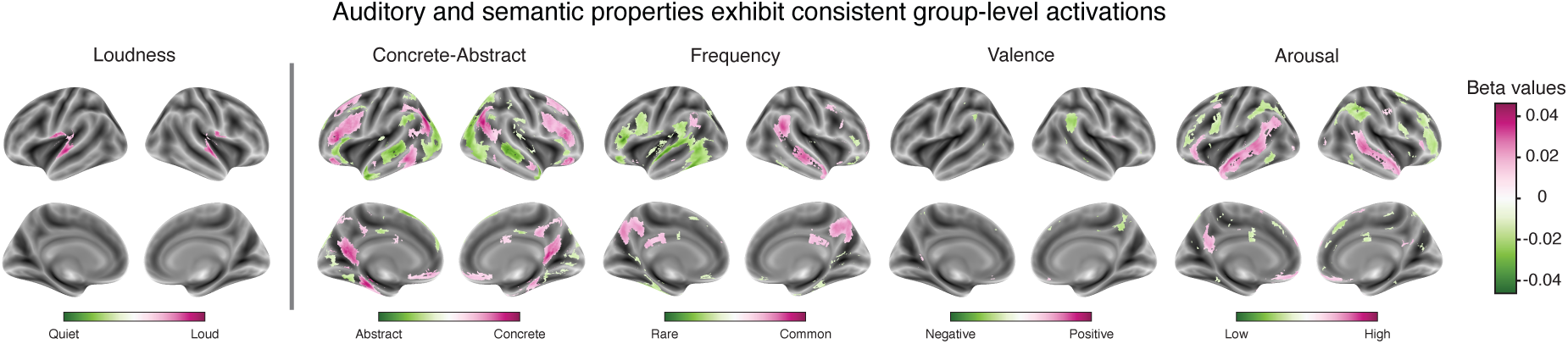
Group-level univariate activation to auditory and semantic properties of language. Across stories and subjects, multiple regions exhibited significant activation to the intensity of sound and word-level semantic properties including concreteness, prevalence, valence, and arousal. Results shown are from a single linear mixed-effects model containing fixed effects for all linguistic properties plus random effects for story and subject. Results are displayed at a voxel-wise false-discovery rate (FDR) threshold of *q* < 0.05.

**Figure 3.**
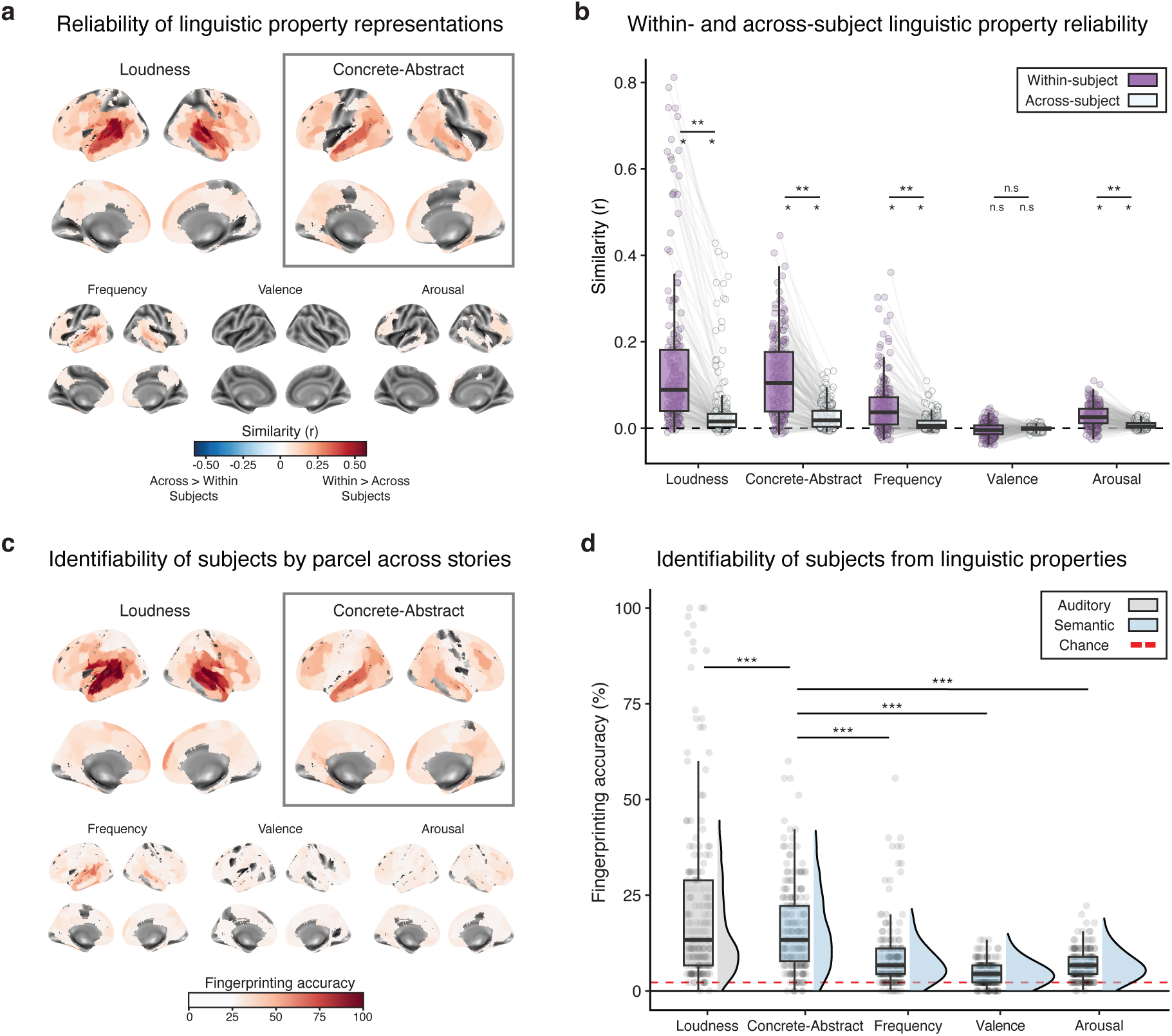
Within- and across-subject reliability of neural representations of linguistic properties. We compared representations of linguistic properties across four naturalistic stories both within and across subjects. **(a)** Across stories, all linguistic properties (excluding valence) exhibited high within-subject reliability across a wide-swatch of cortex (*p* < 0.05, null = 10,000 permutations, corrected for multiple comparisons using FDR correction). **(b)** While a simple auditory property, loudness, exhibited the highest reliability, representations of the concrete-abstract axis were more reliable than other semantic properties (frequency, valence, arousal). Across all linguistic properties, within-subject reliability was consistently higher than across-subject reliability. **(c)** Representations of linguistic properties enabled accurate identification of subjects across a wide swath of cortex. All plots are threshold at chance (2.22%). **(d)** Out of tested semantic properties, subjects were most identifiable from their representations of the concrete-abstract axis. Each dot indicates the reliability within a parcel of the Schaefer parcellation (200 total). * *p* < 0.05; ** *p* < 0.01; *** *p* < 0.001; n.s. *p* > 0.05.

#### Evaluating the reliability of representations of semantic properties

Next, to understand whether semantic properties elicit reliable representations during story listening (Figure 1C), we examined the within- and across-subject multivariate pattern similarity of evoked responses for each property across stories. We divided the cortex into 200 parcels using the Schaefer parcellation^39^. Then, within each parcel, we correlated the voxel-wise beta values across all pairs of participants for all unique pairs of stories (six total pairs) before averaging across all story-pair matrices to obtain a subject-pairwise similarity matrix. We repeated this process for each property to understand the similarity of neural representations across stories both within- and across-subjects. See Figure 1D for a schematic of this analysis.

We evaluated two multivariate signatures of these neural representations (Figure 1D). Our first method — reliability — evaluates the similarity of a subject’s representations to themselves across stories compared to the similarity of their representations to others. Specifically, reliability is calculated as the difference between the similarity of a subject to themselves (within-subject similarity) and the average pairwise similarity of a subject to all other subjects (across-subject similarity).

Our second method — identifiability — measures how unique representations are to each subject. A subject is said to be identifiable based on their representations when, across stories, within-subject similarity is higher than similarity to all other participants of the group. For each parcel, we calculate identifiability as fingerprinting accuracy: the average number of participants identifiable based on their neural representations^51^.

For reliability analyses, statistical significance was evaluated via permutation testing (null = 10,000 permutations). For identifiability analyses, statistical significance was evaluated against chance (2.22%, or 1/45, where 45 is the total number of subjects). Resulting *p*-values for each signature were corrected for multiple-comparisons across 200 parcels using the Benjamini-Hochberg method (*q* < 0.05). To evaluate reliability and identifiability at a whole-brain level, for each signature, we used a linear-mixed effects model to predict reliability/identifiability from the fixed-effect of linguistic property while controlling for the random effect of parcel in both models and a random effect of subject within the reliability model. Then, to test for significant differences between linguistic properties, we conducted pairwise statistical tests between model fits to each property. We also conducted one-sample tests for both the within- and across-subject reliability for each linguistic property. All tests were two-tailed, tested at alpha *p* < 0.05, and corrected for multiple-comparisons using FDR correction.

#### Disentangling the reliability of concrete and abstract word representations

We next aimed to understand whether concrete and abstract words differentially drive reliability of neural representations of the concrete-abstract axis. To this end, within each story, we limited our analysis to nouns (as verbs were more prevalent at the abstract end) and dichotomized the concrete-abstract axis by selecting the top 30% of concrete and top 30% of abstract words (Figure 4A). Specifically, we asked if and where concrete word representations are more reliable than abstract word representations or vice versa.

**Figure 4.**
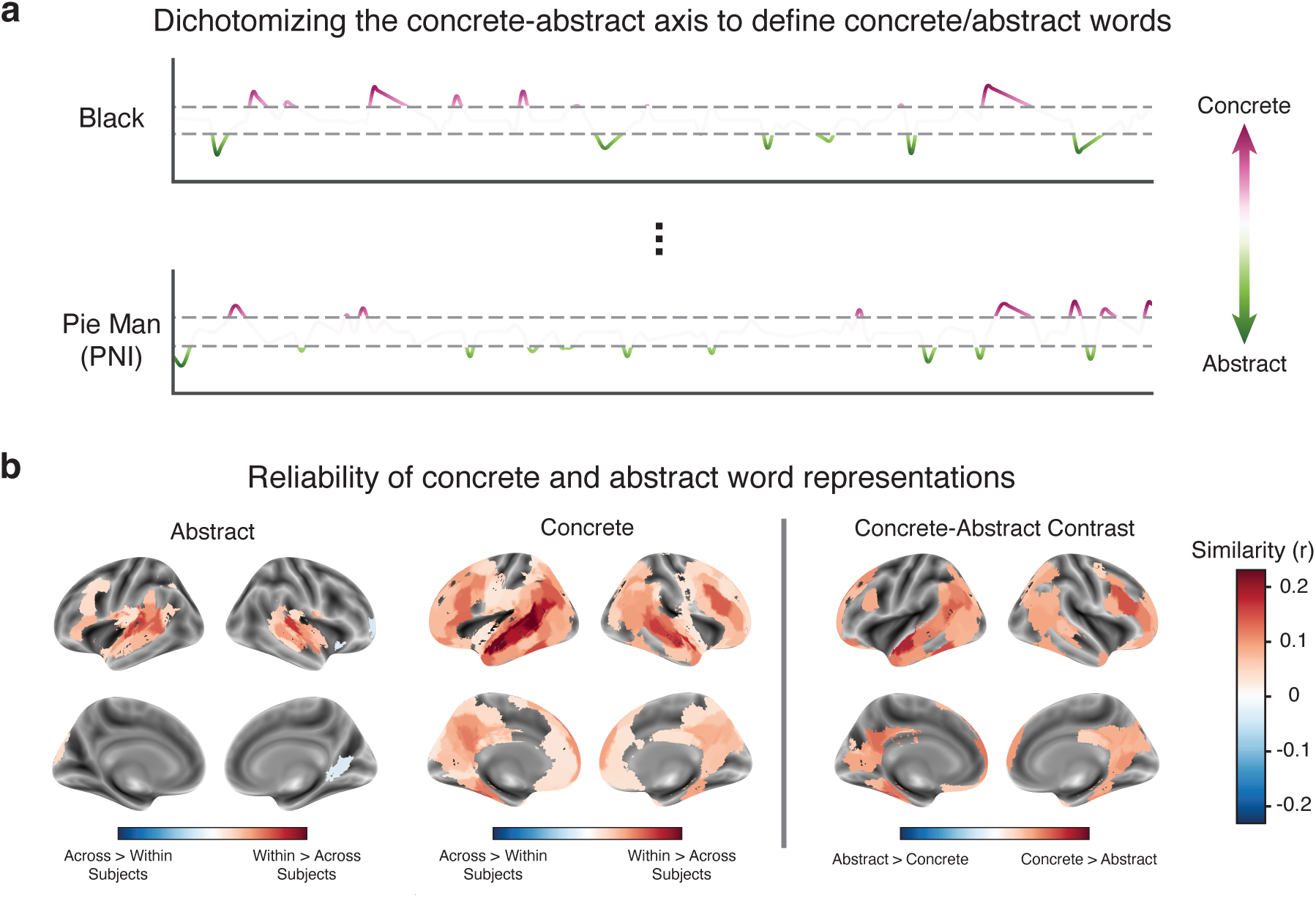
Within-subject reliability of neural representations of concrete and abstract words. **(a)** We selected concrete and abstract words as the top/bottom 30% of the concrete-abstract axis and estimated neural responses to each set of words in a second GLM analysis. **(b)** While both concrete and abstract words exhibited reliable representations within subjects across stories, concrete words were more reliable than abstract words. *p* < 0.05, null = 10,000 permutations.

We used a GLM to estimate separate BOLD response patterns for concrete and abstract words (using regressors defined based on the top 30% of each end). Within this model, we specified concrete and abstract words as event regressors, discarding the amplitude component and treating all words of a given property as contributing equally to the model of BOLD response. We also included two amplitude-modulated regressors, word frequency and loudness, to control for differences in low-level semantic and auditory features. We then repeated our analysis of reliability (described above) on the beta maps of concrete and abstract words.

For each parcel, we contrasted concrete and abstract word reliability within each subject by applying Fisher’s z-transformation and taking the difference between the reliability scores (concrete minus abstract), limiting our analysis to parcels that showed significant reliability for *either* concrete or abstract words. Then, within each parcel, we conducted paired t-tests to identify parcels that significantly differed in their reliability of concrete and abstract word representations. All tests were two-tailed, tested at alpha *p* < 0.05, and corrected for multiple-comparisons using FDR correction.

#### Evaluating the stability of concrete and abstract word representations

In light of the finding that representations of concrete words are more reliable than those of abstract words (cf. Fig. 4), we asked whether this higher reliability is driven by more stable semantic relationships between words at the concrete end of the spectrum. To define semantic relationships between words, we used a natural language processing model (GloVe)^52^ to embed each word in both the top 30% concrete and top 30% abstract word sets, aggregated across stories, within a high-dimensional semantic space (Figure 5A). We then applied spectral clustering^53^ over the concrete and abstract word embeddings to obtain concept clusters for each end of the spectrum (k=3 each for the concrete and abstract ends, so six total) composed of semantically similar words. While we selected k=3 clusters to balance the number of words in each cluster, similar results were obtained at both k=2 and k=4 clusters. These clusters grouped concrete and abstract words into sets of related concepts — such as a food-related concrete cluster containing the words “bread” and “cheese” — that were visually distinct when projected into a 2-dimensional space using UMAP^54^. Importantly, words within each concept cluster could come from within the same story or from different stories.

**Figure 5.**
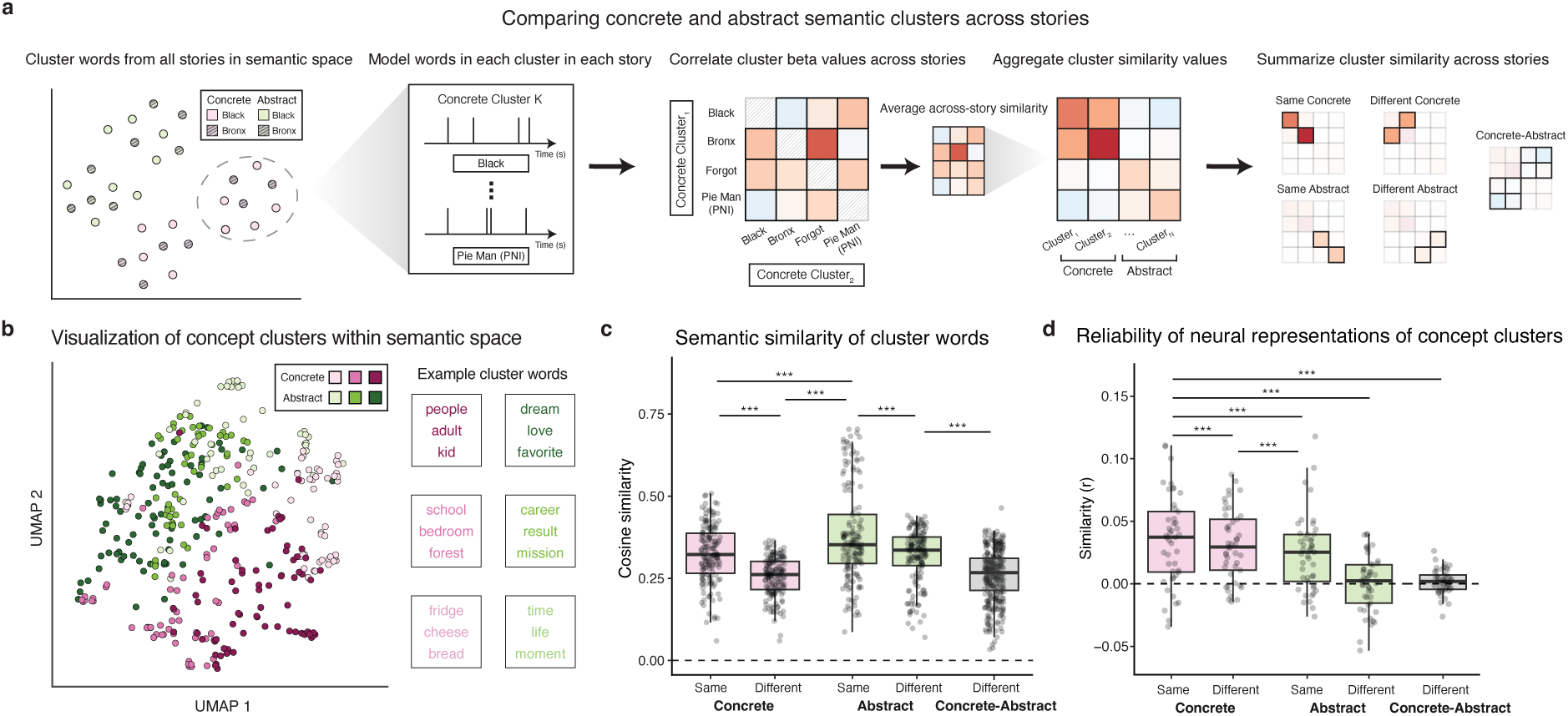
Stability of concrete and abstract word representations within and across subjects. **(a)** We embedded and clustered the top 30% concrete and top 30% abstract words within a high-dimensional semantic space (GloVe). We then estimated voxel-wise beta values for each of six clusters (3 concrete, 3 abstract) within each subject and story. Next, within each parcel (200 total), we correlated beta values between all sets of clusters across stories and averaged the across-story similarity of clusters. **(b)** Visualization of word embedding clusters within a 2-dimensional projection using UMAP. **(c)** Within semantic space, words within abstract clusters were more similar (i.e., less distant) than words within concrete clusters. Each dot represents the average similarity of a given word to other words within a given comparison. In contrast, **(d)** within-subject neural representations of concrete clusters were more similar across stories than representations of abstract clusters. Each dot indicates the average similarity of a subject’s concept cluster representations within a given comparison. * *p* < 0.05; ** *p* < 0.01; *** *p* < 0.001; n.s. *p* > 0.05.

In addition to visualizing the qualitative organization of concept clusters, we also formally tested the semantic similarity of words in the same or in different clusters, within and between ends of the concrete-abstract spectrum. Importantly, because the clustering itself was done on semantic distances, we expect that distances will be lower between words in the same versus different clusters, but this analysis also lets us quantify if and how semantic spread *across* clusters is greater at one end of the concrete-abstract axis than the other. Specifically, we calculated the cosine similarity between all pairs of words embedded within the semantic space. We then grouped these pairwise similarity values into the following categories: a) pairs of words within the same cluster, b) pairs of words in different clusters at the same end of the concrete-abstract axis (i.e., either concrete or abstract), and c) pairs of words at different ends of the concrete-abstract axis, which were (by definition) in different clusters. To compare these groups of similarity values, we used a linear-mixed effects model to evaluate how end of the property spectrum (concrete vs. abstract), cluster membership (within vs. between), and the interaction between these two features relate to the semantic similarity of cluster words while controlling for the random effect of word. To help interpret any resulting differences, we also conducted follow-up pairwise statistical tests. All tests were two-tailed, tested at alpha *p* < 0.05, and corrected for multiple-comparisons using FDR correction.

Next, we used a GLM to estimate BOLD responses to words within each concept cluster and evaluated the similarity of neural concept-cluster representations across stories. Similar to our analysis of semantic space, we calculated a) the similarity of neural representations of the same cluster across stories, b) the similarity of neural representations of different clusters at the same end of the spectrum (e.g., concrete clusters to other concrete clusters), and c) the similarity of neural representations between concrete clusters and abstract clusters. Crucially, all analyses of cluster similarity, both within- and across-subjects, are calculated as the similarity of clusters *across stories*; this allowed us to evaluate the stability and uniqueness of concept-cluster representations across distinct presentations and contexts.

Using two separate linear-mixed effects models, we examined how end of the property spectrum (concrete vs. abstract), cluster membership (within vs. between), and specific cluster relationship (e.g., within-concrete, between-concrete, etc.) differentially contribute to whole-brain similarity of neural representations while controlling for random effects of subject and parcel. Our first model predicts similarity from the fixed-effects of end of the property spectrum and cluster membership, and evaluates their main effects as well as their interaction. Then, in a separate model, we predict similarity from the fixed-effect of specific cluster relationship, specifying each cluster relationship as a separate level of the fixed effect. Using this second model, we tested for significant differences between cluster relationships by conducting pairwise statistical tests. All tests were two-tailed, tested at alpha *p* < 0.05, and corrected for multiple-comparisons using FDR correction.

## Results

We aimed to understand how neural representations of the concrete-abstract axis vary within individuals and across the population during naturalistic story listening. Using a large dataset of subjects (N=45) that listened to four stories each, we replicated previous findings that neural responses to the concrete-abstract axis show group-level consistency. Complementing this consistency, we also found idiosyncratic representations that were unique to individuals and stable across stories, allowing us to identify subjects with a high degree of accuracy. Furthermore, by placing words within a high-dimensional semantic space, we demonstrated that neural representations of concrete words are particularly stable and stereotyped, and that this consistency primarily drives the reliability of the concrete-abstract axis, while representations of abstract words are more variable both within and across subjects.

### Consistent group-level representations of the concrete-abstract axis

We first sought to replicate prior work that demonstrates group-level consistency of univariate activity to concrete and abstract words. For each subject and story, we modeled brain activity as a function of the time-varying concreteness level of its content (as given by word-level norms provided by a separate set of human raters). Our model also included time-varying regressors for other semantic properties — namely, frequency, valence, and arousal — plus loudness, a low-level auditory control.

All linguistic properties demonstrated neural responses consistent across both subjects and stories (Figure 2; *q* < 0.05). For example, loudness evoked responses in bilateral primary auditory cortex. Critically, the concrete-abstract axis evoked neural responses across a wide swath of cortex: more concrete words drove higher responses in regions including bilateral angular gyrus, bilateral parahippocampal cortex, and bilateral inferior frontal gyrus, while more abstract words drove responses in regions such as bilateral superior temporal gyrus and bilateral anterior temporal lobe. These results align with previous research that has reported similar cortical regions engaged in processing concrete and abstract concepts^55,56^. Importantly, all semantic properties exhibit responses that replicate prior research on word property representation: frequency modulation in the left inferior frontal gyrus^47^, valence representations in the right temporoparietal junction^57^, and arousal representations in posterior cingulate^58^ and ventromedial prefrontal cortex^49^.

### Representations of the concrete-abstract axis are individually reliable

Having shown that the concrete-abstract axis evokes consistent univariate activity at the group level, we next investigated the individual reliability of multivariate representations of this axis as well as other linguistic properties. We found that representations of all properties, excluding valence, exhibited within-subject reliability across stories in at least some brain regions (Figure 3A; n = 10,000 permutations, *p* < 0.001). Importantly, while loudness showed the highest average reliability across parcels (*r* = 0.11) compared to the concrete-abstract axis (*r* = 0.09; β = 0.06, *t*(42967) = 44.75, *p* < 0.001), the concrete-abstract axis showed the second highest average reliability and was significantly more reliable than all other semantic (i.e., non-primary-sensory) properties (frequency: *r* = 0.04, β = 0.01, *t*(42967) = 8.71; valence: *r* = −0.002, β = 0.05, *t*(42967) = 41.83; arousal: *r* = 0.02, β = 0.03, *t*(42967) = 23.74; all *p*s < 0.001).

We next disentangled the separate contributions of within- and across-subject similarity in driving reliability of individual representations. In theory, high individual reliability of representations across stories could result from 1) highly *similar* representations within subjects, 2) highly *dissimilar* representations across subjects, or 3) a combination of the two. Accordingly, for each linguistic property, we computed and compared within- and across-subject similarity of representations. Across all properties with significant reliability (all linguistic properties excluding valence), subjects were significantly similar to themselves (Figure 3B; one-sample t-tests, all *p*s < 0.001) and significantly more similar to themselves than to other subjects (paired t-tests, all *p*s < 0.001). Interestingly, by correlating within- and across-subject similarity values across parcels, we found that, at a whole-brain level, linguistic properties that demonstrated higher within-subject similarity also showed higher across-subject similarity (loudness (*r* = 0.874), concrete-abstract (*r* = 0.784), frequency (*r* = 0.797), valence (*r* = 0.428), arousal (*r* = 0.599); all *p*s < 0.001). This finding recapitulates a seemingly paradoxical phenomenon of individual differences research previously shown in functional connectivity fingerprinting: brain states that make individuals more similar to themselves also make them more similar to others^59^.

### Individuals are identifiable from their representations of the concrete-abstract axis

The previous analysis revealed that individuals’ representations of the concrete-abstract axis are reliable, but how *unique* are these representations? High reliability does not necessarily imply uniqueness: low average across-subject similarity could be due to high *variability* in across-subject similarity. Specifically, select pairs of subjects may possess highly similar representations of the concrete-abstract axis, despite most of the group exhibiting low similarity. To test the extent to which linguistic property representations are unique to each individual, we evaluated our ability to identify subjects from their representations of each word property.

Across cortical parcels, we were able to identify subjects from representations of both sensory response (loudness) and all four semantic properties across much of the brain (Figure 3C; null = 10,000 permutations, all *p*s < 0.001). Of note, the average identification rates across cortical parcels were low in an absolute sense but still significantly above chance (chance = 2.22%; Figure 3D). Overall, representations of loudness provided the best ability to identify subjects (22.1%), demonstrating significantly higher identification rates, on average, than the concrete-abstract axis (16.5%; β = 10.41, *t*(948) = 14.77, *p* < 0.001). Importantly, representations of the concrete-abstract axis enabled significantly higher identification accuracy than representations of other semantic properties (frequency: 8.8%, β = 2.9, *t*(948) = 4.11; valence: 4.4%, β = 7.24, *t*(948) = 10.27; arousal: 6.6%, β = 5.08, *t*(948) = 7.2; all *p*s < 0.001). We then applied a winner-takes-all approach to identifiability maps to understand the cortical parcels where concrete-abstract axis representations showed the highest accuracy out of all linguistic properties. We found that the concrete-abstract axis enabled the highest identification of subjects—even higher than loudness—within regions including left anterior temporal lobe, left inferior frontal gyrus, and bilateral retrosplenial cortex (RSC). These regions dovetail with previous studies that have shown that left-lateralized language network and default mode network (DMN) are important in representing concrete and abstract concepts^28,56,60–63^.

### Concrete word representations are more reliable than abstract word representations

Thus far, we have shown that representations of the concrete-abstract axis are reliable within and unique to individual subjects across experiences. Yet it remains unclear whether both concrete and abstract words contribute equally to driving the reliability of representations. On one hand, concrete words may be more reliable than abstract words because they are less sensitive to the surrounding situational context. On the other hand, abstract words may be more idiosyncratic than concrete words, as uniquely language-based representations could depend more heavily on individual experience to create meaning. While a previous study observed that representations of abstract words exhibited lower similarity across subjects than concrete words, disentangling the source of these results necessitates 1) presenting words within naturalistic contexts and 2) evaluating similarity within subjects, across experiences. To understand the differential contributions of concrete and abstract words in driving reliability, we dichotomized the continuous, concrete-abstract axis into concrete and abstract words and estimated reliability separately for each end of the spectrum.

We observed that representations of concrete and abstract words each demonstrated significant reliability across stories in several brain regions (Figure 4B; null = 10,000 permutations, both *p*s < 0.001). Contrasting the reliability maps for concrete and abstract words, we found that a large number of cortical parcels (36% or 72/200) exhibited more reliable responses to concrete words than abstract words. On the other hand, no parcels showed greater reliability of abstract word representations compared with concrete word representations. This finding suggests that concrete word representations primarily drive reliable responses of the concrete-abstract axis and extends previous, population-level findings to individual neural responses^16,17,28,32,62^.

### Stable clusters of concrete words drive reliability of representations across experiences

Why might neural representations at the concrete end of the spectrum be more reliable than representations at the abstract end? While the naturalistic nature of these stimuli means that we did not necessarily have repeated presentation of the *same* word(s) across stories, we can use natural language processing (NLP) techniques to group words into clusters of semantically related words and use the clusters to help understand why concrete representations are more reliable, even when generalizing over individual words and concepts. Numerous recent studies have demonstrated parallels in language representation between NLP models and human neural processing^13,64–67^. Here, we used a word-embedding NLP model (GloVe)^52^ to understand how the semantic relationships among concrete and abstract words relate to the reliability of their neural representations. Specifically, we embedded concrete and abstract words within a high-dimensional semantic space and clustered words based on their semantic similarity. We then analyzed the similarity of word clusters in semantic space and, separately, the similarity of neural responses to each word cluster across stories using linear mixed-effects models (see Methods).

Before evaluating neural responses to concrete and abstract word clusters, we first examined the similarity of cluster words within the semantic space. Unsurprisingly, words within the same cluster were more similar to each other than to words in different clusters (Figure 5C; β = 0.03, *t*(610) = 14.71, *p* < 0.001), a pattern of results consistent across both concrete and abstract words (pairwise comparisons; concrete: *t*(306) = 10.76; abstract: *t*(306) = 10.03; both *p*s < 0.001). But we also observed a somewhat puzzling result: within semantic space, abstract words were generally more similar to one another than concrete words were to one another (β = 0.03, *t*(610) = 5.87, *p* < 0.001). This finding was particularly surprising given the results from the previous analysis (cf. Fig 4B) that showed neural representations of concrete words to be more reliable than representations of abstract words. Why might the concrete end of the spectrum, which encompasses *more* variability in (i.e., spans more of) semantic space, show *less* variability in its neural representations?

We next turned to analyze within-subject neural representations of concrete and abstract word clusters. Similar to the results in semantic space, representations of words within the same cluster were more similar across stories than representations of words in different clusters (Figure 5D; β = 0.007, *t*(34373) = 20.04, *p* < 0.001), and this was true for both the concrete and abstract ends of the spectrum (concrete *z* = 4.36, abstract *z* = 23.99, both *p*s < 0.001). In contrast to the similarity of clusters in semantic space (Figure 5C), neural representations of concrete words exhibited greater similarity regardless of semantic distance (same or different clusters) than abstract words (β = 0.01, *t*(34373) = 29.45, *p* < 0.001). Critically, there was also an interaction such that the similarity advantage for same-over different-cluster representation was smaller for concrete than for abstract words (β = −0.005, *t*(34373) = −13.88, *p* < 0.001). Strikingly, neural representations of *different* concrete clusters were more similar than neural representations of the *same* abstract cluster (mean difference = 0.007, *z* = 7.12, *p* < 0.001). Furthermore, this pattern of results persisted when analyzing similarity *across* subjects (within > across: β = 0.002, *t*(34373) = 24.11; concrete > abstract: β = 0.001, *t*(34373) = 17.07; interaction: β = −0.001, *t*(34373) = −13.27; all *p*s < 0.001), suggesting that a consistent principle drives the organization of concrete and abstract neural representations both within individuals and across the population.

Considered together, neural representations of distinct concrete concepts were more similar than those of distinct abstract concepts, despite concrete words spanning greater distances within semantic space than abstract words. These divergent results between the NLP model and neural data suggest that concrete words share additional properties beyond purely linguistic representations, such as imageability, that could stem from integrating visual information into the neural representations.

## Discussion

Word meanings vary both across people and contexts, often informed by both conceptual associations specific to the individual and different situations in which the word is used. What semantic properties enable convergent conceptual knowledge while simultaneously supporting unique, individual experience? Here, we found that the concrete-abstract axis provides a basis for both population stability and individual variability in representation of natural language.

Our results provide further evidence for the importance of the concrete-abstract axis in semantic representations of language. Numerous studies have demonstrated that, while both concrete and abstract words evoke responses within the language network^28,30,68,69^, responses to concrete words are generally stronger and longer-lasting than responses to abstract words^16,31,32,70^. Prior work has also shown that concrete words engage areas beyond the language network, such as the default mode network (DMN), more than abstract words^28,56,60,61,63^. In our study, we found reliable representations of the concrete-abstract axis within both the language network and DMN that were unique to individual subjects across diverse, naturalistic stories. While an auditory property — loudness — exhibited the most reliable representations across stories, it is likely that this property contained additional language-related information beyond pure audition due to the presence of few other semantic properties. Critically, representations of the concrete-abstract axis were more reliable than representations of other semantic axes (i.e., frequency, valence, arousal), driven primarily by the reliability of concrete word representations (as opposed to abstract word representations). Together, our results suggest that the reliability of concrete word representations may be due to engagement of areas beyond the language network, including DMN, that engage more imagery-related processes than abstract words and other semantic properties.

Traditionally, neural representations of language have been probed by presenting participants with single words, sentences, and short paragraphs^71,72^. These studies have revealed neural territory specific to language^5,73^ that closely interacts with other networks involved in cognitive control and theory of mind^4,74,75^. In contrast to these carefully controlled experiments, everyday language is dynamic and contextualized – the meanings of words and sentences are informed by larger narrative structure^33,76^. It is therefore crucial to evaluate the degree to which findings of carefully-controlled studies extend to naturalistic language perception^77^. Within the present study, participants were presented with naturalistic auditory narratives representative of how language is used in day-to-day life. Importantly, we found that representations of abstract words were more variable both within and across subjects than representations of concrete words.

The finding of higher across-subject variability for abstract words aligns with another recent study that used a single-word paradigm^17^; the authors of that study interpreted this heightened variability as reflecting individual differences in meaning of abstract words in particular. However, the appeal to individual differences implies a stability of representations *within* the same subject over time, which was not tested. Our study differs from this previous work in two ways: first, we examined word and concept representations within subjects across repeated presentations, and second, we captured these neural representations during a naturalistic listening task that presented words in context. We found that compared to representations of concrete words, representations of abstract words and concepts were not only more variable across subjects, but also within the same individual across distinct experiences. This suggests that variability in abstract words stems less from individual differences in meaning and more from a general instability of their representations, perhaps because their meanings are more context-dependent.

Recent developments in natural language processing (NLP) models have provided researchers with tools to better investigate how the human brain organizes and processes natural language^13,64–67^. These computational models not only capture semantic relationships between words, but also contain rich knowledge regarding how words relate within various contexts^78^. Importantly, the contextual relationships between concrete words — that a fish and a whale may be semantically similar in terms of “wetness” but different in terms of “size” — closely correspond to human judgements of the same categories^79^. Yet, within our study, we found that clusters of concrete words were less similar than clusters of abstract words within an NLP model but *more* similar in the human brain. This dissociation suggests that neural representations of the concrete-abstract axis contain additional information beyond pure linguistic representation. Given the close relationship between concreteness and imageability^21,22^, concrete words may carry a signature of imageability that results from being jointly represented across visual and linguistic domains, thereby boosting the stability of their neural representations both within and across subjects.

Though our work aligns with and extends past work on the concrete-abstract access, we highlight the following limitations. First, it is possible that neural representations of other semantic axes are also idiosyncratic. In the current study, we specifically leveraged human ratings of words along semantic axes, but these behavioral ratings were collected by presenting participants with individual words out of context. Similarly, we leveraged an NLP model that does not incorporate contextual information into the word-level representations. Other semantic axes, such as valence and arousal, may be more context-dependent and require ratings specific to a given story or individual to understand the idiosyncrasies in neural representations. Second, due to the diversity of content across the auditory narratives, we were limited in our ability to compare representations of the *same* words across stories. We addressed this by comparing the neural representations of clusters of similar words across stories, extending prior work on single words to the organization of broader concepts in semantic space. Future work could select stories that contain the same words but vary in narrative content to understand the stability of both specific words and semantic organization more generally across experiences.

In sum, our work establishes the concrete-abstract axis as a critical dimension for promoting both shared and individualized representations of language. In particular, these findings disentangle the sources of individual variability of concrete and abstract concept representations. Our results underscore the importance of considering within-subject variability in the context of the broader population to differentiate underlying drivers of idiosyncratic processing of natural language.

